# Improved coral thermal tolerance through modulation of antioxidant defenses

**DOI:** 10.64898/2026.05.21.723506

**Authors:** Laura F.B. Marangoni, Eric Beraud, Maria Chacon, Oren Levy, Miguel Mies, Matthieu Leray, Christine Ferrier-Pagès

## Abstract

Mass coral bleaching events, driven by rising ocean temperatures, are pushing reef ecosystems toward collapse on a global scale. Because oxidative stress is an early driver of coral bleaching, strategies that enhance coral antioxidant defenses may improve coral resilience under thermal stress. Here, we tested a targeted antioxidant supplementation designed to enhance oxidative stress regulation in three representative Red Sea scleractinian coral species subjected to a thermal challenge. While responses varied among species and physiological metrics, supplemented corals consistently maintained higher photophysiological performance under heat stress. In *Stylophora pistillata*, antioxidant supplementation was associated with enhanced catalase activity, maintenance of glutathione redox homeostasis, and lower intracellular reactive oxygen species levels. In contrast, non-fed corals exhibited oxidative imbalance, increased lipid peroxidation, and impaired photophysiological recovery, while corals receiving a non-enriched heterotrophic diet showed an intermediate response characterized by increased catalase activity but persistent glutathione oxidation and elevated ROS during recovery. Together, the dietary treatments revealed a gradient in oxidative regulation, ranging from insufficient antioxidant protection in autotrophic corals to enhanced oxidative homeostasis in antioxidant-supplemented corals. Our findings demonstrate the potential of targeted nutritional antioxidant supplementation to enhance coral oxidative regulation and physiological performance under elevated temperatures, highlighting a promising complementary approach for coral conservation and restoration efforts.

## Introduction

Global temperatures have risen by approximately 1.5 °C since the onset of the Industrial Era due to anthropogenic greenhouse gas emissions ^1^. As a result, live coral cover is projected to decline to less than 10% of 1990s levels ^2,3,4^. Ocean warming is indeed the primary cause of mass coral bleaching events, during which reef-building corals lose their symbiotic dinoflagellates (family *Symbiodiniaceae*) under thermal stress ^5^. The loss of these symbionts severely compromises coral physiology, leading to reduced energetic acquisition, lower calcification rates, and increased susceptibility to disease and mortality. Repeated or prolonged bleaching events, such as those observed during recent global marine heatwaves, have led to widespread degradation of reef ecosystems, with significant losses in biodiversity and ecosystem function ^6,7,8^.

Coral bleaching results from a complex set of processes, and different hypotheses have been proposed to explain the breakdown of the coral-algae symbiosis, including the carbon limitation and the oxidative stress hypotheses ^9,10^. The carbon limitation hypothesis suggests that stress disrupts the transfer of photosynthetically fixed carbon from the symbionts to the coral host, creating an energetic imbalance ^11,12^. As the autotrophic supply of nutrients is compromised, corals need to depend more on heterotrophic intake (capture of planktonic prey) and energy reserves when possible ^13,14,15^. Heterotrophy therefore plays a key role in coral resilience to bleaching by facilitating recovery of symbiotic function after thermal stress ^16^; providing lipids for gamete production during reproduction, even under severe bleaching ^17^; and supporting juvenile growth during the vulnerable post-settlement stages ^18^; and post-transplant survival ^19^. Importantly, it also supplies essential metals (e.g., Mn, Fe, Mg) that are cofactors for key antioxidant enzymes ^20,21,22^. The oxidative stress hypothesis posits that thermal stress disrupts the photosynthetic apparatus of the symbiotic algae, leading to the accumulation of reactive oxygen and nitrogen species (ROS/RNS) that trigger breakdown of the symbiosis ^9,10^.

High ROS levels indeed disrupt cellular metabolic and signaling pathways and lead to oxidative damage in macromolecules (e.g., lipids, proteins, and DNA), which, if unbalanced, can result in cell death ^23,24^. To cope with ROS, corals rely on both low molecular weight antioxidants and antioxidant enzymes ^9^. Antioxidant enzymes [*e.g*., superoxide dismutase (SOD), catalase (CAT), and peroxidases] are often considered the most important system because they can catalytically neutralize large amounts of ROS and provide a sustainable and energy-efficient defense ^23,25^. Low molecular weight antioxidants, such as glutathione and vitamins E, A, and C, among others ^26,27^, also play essential roles as secondary defense mechanisms, although their reserves can be rapidly depleted ^25^. The interplay between these antioxidant mechanisms is crucial for the stress response of corals.

The carbon limitation and oxidative stress frameworks are not mutually exclusive. Rather, evidence supports an integrated model in which oxidative stress and metabolic imbalance jointly contribute to bleaching outcomes, among other processes ^10,28^. Boosting coral antioxidant capacity may therefore represent a promising avenue to increase stress tolerance ^29^, with nutritional processes, including heterotrophic feeding, playing a central role in this strategy. Boosting coral antioxidant capacity may therefore represent a promising avenue to increase stress tolerance ^29^, with nutritional processes, including heterotrophic feeding, playing a central role in this strategy.

Inspired by the concept of “superfoods” in human nutrition—nutrient-enriched formulations designed to enhance natural defenses and promote overall well-being ^30,31^—we formulated a coral-specific ‘superfood’ aimed at fortifying antioxidant defenses. Heterotrophic feeding through the provision of zooplankton such as *Artemia salina* nauplii is already a well-established pathway to enhance coral resilience by supplementing energy reserves, building tissue biomass, and supporting symbiont recovery during stress. A preliminary dietary intervention in which *Artemia* prey were enriched in lipids was shown to improve coral performance under thermal stress ^32^, highlighting the potential of targeted supplementation to improve coral resilience. In addition, exogenous antioxidants have shown promise in mitigating bleaching, whether delivered directly in seawater or embedded in gels ^33,34^. However, these approaches are limited by rapid photodegradation, which reduces their effectiveness ^35^. Incorporating antioxidants and enzymatic cofactors into Artemia prey, therefore, represents a more effective strategy, enabling controlled delivery while leveraging a natural feeding pathway. Building on this concept, we developed a more comprehensive supplemental diet formulation, focusing not on lipids but on antioxidants and enzymatic cofactors aimed at enhancing coral oxidative stress responses.

Here, we evaluate the efficacy of a supplemental diet formulation, tailored to the unique physiology of scleractinian corals, as a strategy to enhance their antioxidant defenses and improve their resistance to thermal stress. This approach aims to provide a tool for localized interventions to support reef resilience ^7, 36^. Although coral reef restoration efforts have expanded in recent decades, their long-term success remains constrained by recurrent marine heatwaves, which drive widespread bleaching and mortality in both wild and outplanted corals ^37,38,39^. It is increasingly evident that restoration outcomes are improved when transplanted corals are preconditioned to better withstand environmental stress ^10,40^, highlighting the importance of enhancing coral stress tolerance as a key research priority ^6,40^. To test this concept, we conducted two controlled experiments on three coral species representative of the Red Sea - *Turbinaria reniformis*, *Pocillopora damicornis*, and *Stylophora pistillata* – under different dietary nutritional regimes and thermal stress conditions. Using *S. pistillata* as a model organism, we further investigated biochemical markers of oxidative stress and antioxidant activity. Our findings demonstrate that boosting antioxidant capacity through diet can support coral physiological performance under thermal stress and provide a biologically grounded approach to complement reef conservation and restoration efforts.

## Results

### 1. Experiment 1 - Effects of diet and heat stress on *Turbinaria reniformis* and *Pocillopora damicornis*

Corals were exposed to one of three feeding treatments before thermal stress: an autotrophic diet (Non-Fed), where corals relied solely on photosynthesis; a mixotrophic diet (Fed) using non-enriched *Artemia salina*; and an antioxidant-enriched diet (SuperFed) using *Artemia* supplemented with antioxidant compounds and enzymatic cofactors. After two weeks, half of the corals in each feeding group were exposed to elevated temperature (30 °C for 7 days), and physiological traits were assessed.

#### 1.1) Antioxidant supplementation enhances specific physiological traits under control temperature

At 26 °C, feeding treatment did not affect tissue parameters (symbiont density, chlorophyll concentration, or protein content) in *T. reniformis* (Fig. 1a–d). However, photosynthetic efficiency (ETRmax) was significantly higher in SuperFed corals compared to the other diet groups (Tukey test, p = 0.007; Fig. 1e).

**Figure 1.**
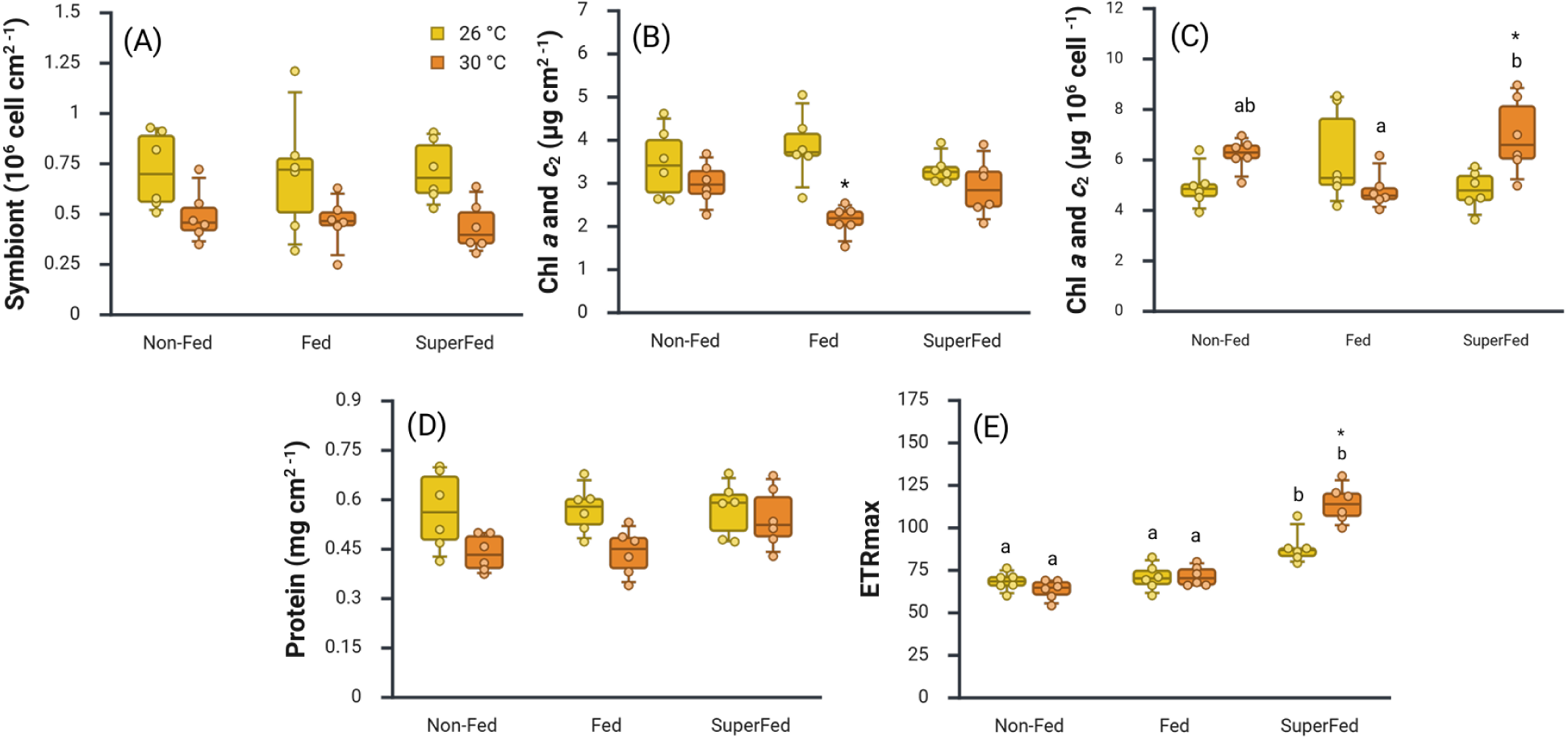
Effect of diet and increased temperature on the physiological response of *Turbinaria reniformis*. Corals were maintained under three different diets [(i) Non-Fed (autotrophic diet - where corals were not fed), (ii) Fed (mixotrophic diet - where corals were fed *Artemia salina* metanauplii, and (iii) SuperFed (superfood diet – where corals were fed with enriched *Artemia salina* metanauplii)], and two thermal conditions of 26 °C (control) and 30 °C (thermal stress): (a) symbiont density, (b) total chlorophyll concentration per surface area, (c) total chlorophyll concentration per symbiont cell, (d) host protein content, and (e) maximum electron transport rate (ETRmax). Data are presented as box plots showing the median, interquartile range, and individual data points. Different letters indicate significant differences (p ≤ 0.05) among dietary treatments at the same temperature, while asterisks indicate significant differences (p ≤ 0.05) between temperatures within the same diet treatment.

In *P. damicornis*, the SuperFed group showed reduced symbiont density (Tukey test, p = 0.012; Fig. 2a). ETRmax was also significantly higher in both Fed and SuperFed corals compared to Non-Fed corals (Tukey test, p < 0.0001; Fig. 2e), while chlorophyll concentration per surface area and symbiont cell, as well as host protein levels, remained unchanged (Fig. 2b, d).

**Figure 2.**
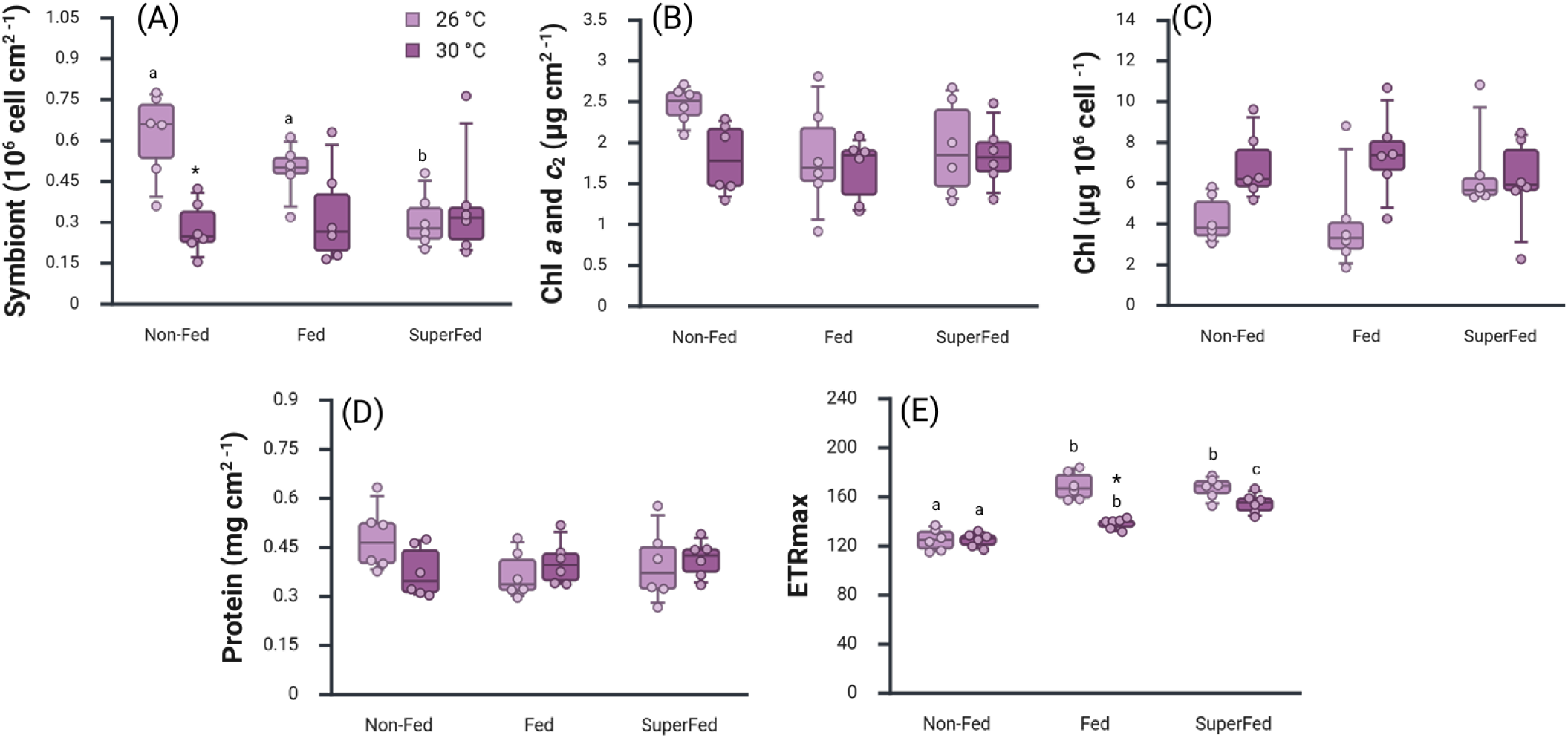
Effect of diet and increasing temperature on the physiological response of *Pocillopora damicornis.* Corals were maintained under three different diets [(i) Non-Fed (autotrophic diet - where corals were not fed), (ii) Fed (mixotrophic diet - where corals were fed *Artemia salina* metanauplii, and (iii) SuperFed (superfood diet – where corals were fed with enriched *Artemia salina* metanauplii)] and two thermal conditions of 26 °C (control) and 30 °C (thermal stress): (a) symbiont density, (b) total chlorophyll concentration per surface area, (c) total chlorophyll concentration per symbiont cell, (d) host protein content, and (e) maximum electron transport rate (ETRmax). Data are presented as box plots showing the median, interquartile range, and individual data points. Different letters indicate significant differences (p ≤ 0.05) among dietary treatments at the same temperature, while asterisks indicate significant differences (p ≤ 0.05) between temperatures within the same dietary treatment.

#### 1.2) Species- and diet-specific physiological responses to thermal stress, with benefits from antioxidant supplementation

At 30°C, physiological responses varied across diets and species. In *T. reniformis*, Fed corals were the only treatment showing a decrease in chlorophyll concentration per surface area relative to their respective control condition (Tukey test, *p* < 0.001; Fig. 1b). SuperFed corals increased their chlorophyll concentration per symbiont cell with respect to their control and to Fed corals at 30 °C (Tukey test, *p* < 0.035; Fig. 1b). Additionally, SuperFed corals exhibited the highest ETRmax at both 26°C and 30°C (Tukey test, *p* < 0.001; Fig. 1e). No changes in symbiont density or in protein content were observed (Fig. 1a and d).

In *P. damicornis*, symbiont density declined significantly under thermal stress in Non-Fed corals (Tukey test, *p* = 0.005). Fed corals showed reduced ETRmax (Tukey test, *p* < 0.013; Fig. 2e), which, however, remained higher than Non-fed corals (Tukey test, p < 0.001). In contrast, SuperFed corals remained largely stable, with no significant change in symbiont traits or in ETRmax, which remained higher than the ETRmax of both Non-fed and Fed corals at 30°C (Tukey test, p < 0.019). While SuperFed corals exhibited the highest ETRmax among treatments, Non-Fed corals showed the lowest values (Fig. 2e). No changes in chlorophyll concentration or in protein content were observed (Fig. 2b,c and d).

Details of 2-way factorial ANOVA are available in Table S1 and Tuke’s multiple comparison test summary in Table S4.

### 2. Experiment 2 - Effects of diet and thermal stress in *S. pistillata*

#### 2.1) Physiological traits and oxidative metabolism under control temperature

In Phase 2, Non-Fed corals showed the highest chlorophyll concentration per symbiont cell, and higher chlorophyll concentration per skeletal area compared to Fed corals (Tukey, p < 0.04, Fig. 3b and c). SuperFed corals exhibited higher symbiont density than Fed corals in Phase 3 (Tukey, p < 0.005, Fig. 5a), and Non-Fed corals showed higher ETRmax than Fed corals (Tukey, p < 0.005, Fig. 5e). Protein content and oxidative stress markers did not differ among diets under control conditions (Figs. 3d and 4).

**Figure 3.**
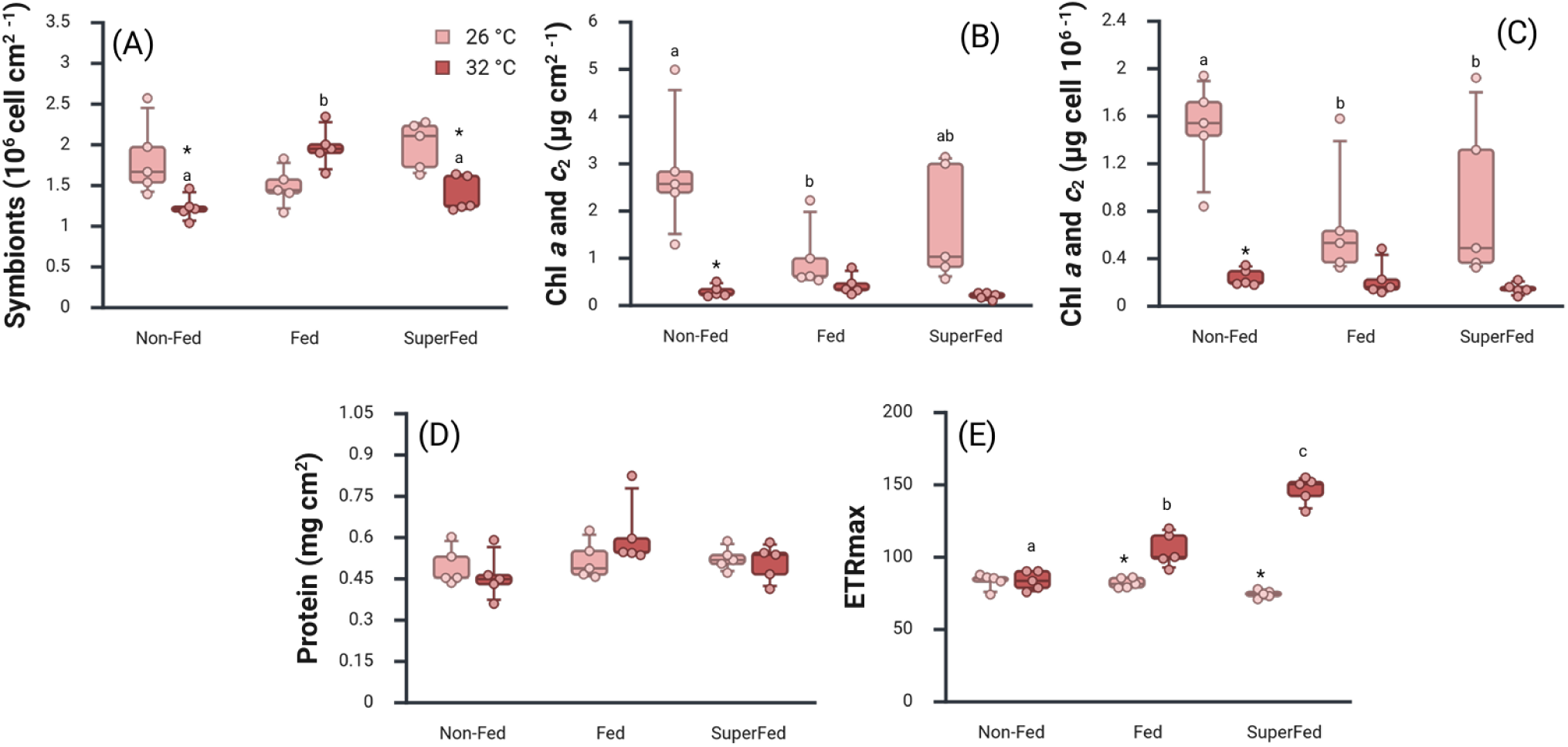
Effect of diet and increasing temperature on the physiological response of *Stylophora pistillata.* Corals were maintained under three different diets [(i) Non-Fed (autotrophic diet - where corals were not fed), (ii) Fed (mixotrophic diet - where corals were fed *Artemia salina* metanauplii, and (iii) SuperFed (superfood diet – where corals were fed with enriched *Artemia salina* metanauplii)] and two thermal conditions of 26 °C (control) and 30 °C (thermal stress): (a) symbiont density, (b) total chlorophyll concentration per surface area, (c) total chlorophyll concentration per symbiont cell, (d) host protein content, and (e) maximum electron transport rate (ETRmax). Data are presented as box plots showing the median, interquartile range, and individual data points. Different letters indicate significant differences (p ≤ 0.05) among dietary treatments at the same temperature, while asterisks indicate significant differences (p ≤ 0.05) between temperatures within the same dietary treatment.

**Figure 4.**
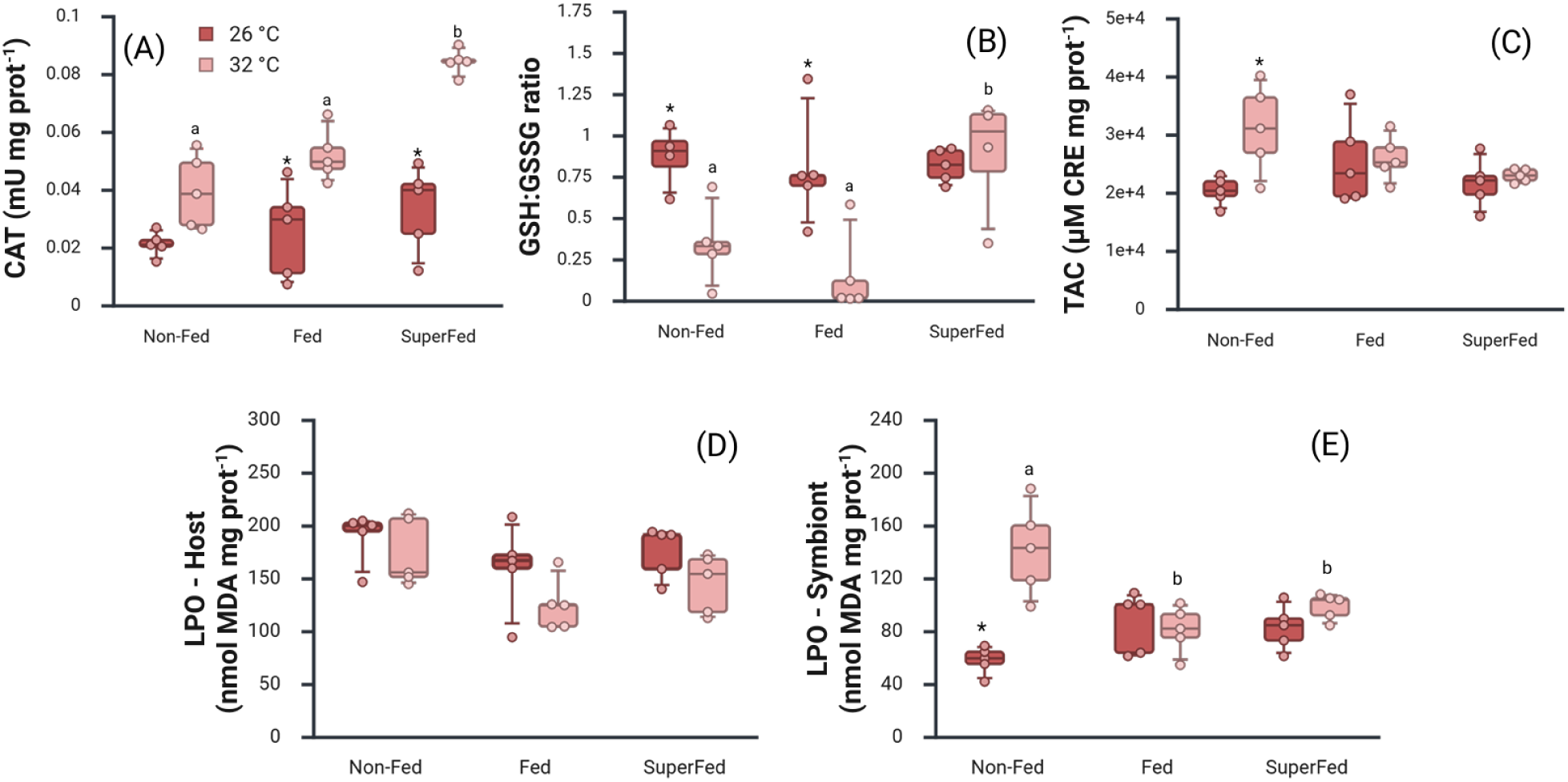
Effect of diet and increased temperature on the oxidative metabolism of *Stylophora pistillata*. (a) Catalase activity, (b) reduced and oxidized glutathione ratio (GSH/GSSG), (c) non-enzymatic total antioxidant capacity (TAC), and oxidative damage to lipids (LPO) in (d) the coral host, and (e) symbionts for *Stylophora pistillata* under three different diets [(i) Non-Fed (autotrophic diet - where corals were not fed), (ii) Fed (mixotrophic diet - where corals were fed *Artemia salina* metanauplii, and (iii) SuperFed (superfood diet – where corals were fed with enriched *Artemia salina* metanauplii)] and two thermal conditions of 26 °C (control) and 30 °C (thermal stress).). Data are presented as box plots showing the median, interquartile range, and individual data points. Different letters indicate significant differences (p ≤ 0.05) among dietary treatments at the same temperature, while asterisks indicate significant differences (p ≤ 0.05) between temperatures within the same diet treatment.

**Figure 5.**
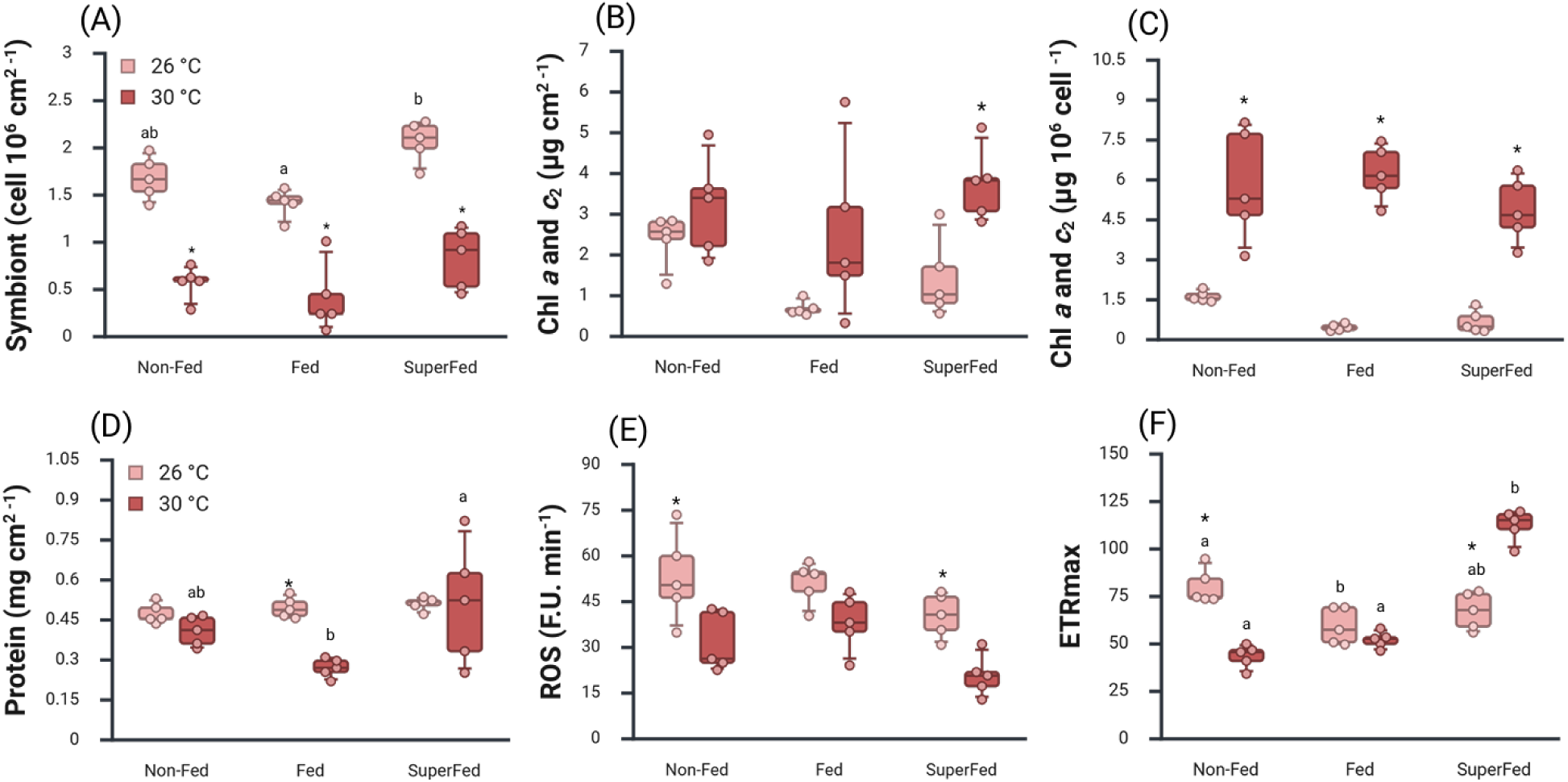
Effect of diet on the recovery of *Stylophora pistillata*. (a) Symbiont density, (b) total chlorophyll concentration per surface area, (c) total chlorophyll concentration per symbiont cell, (d) host protein content, (e) reactive oxygen species (ROS), and (f) maximum electron transport rate (ETRmax) under three dietary treatments: (i) Non-Fed (autotrophic diet - where corals were not fed), (ii) Fed (mixotrophic diet - where corals were fed *Artemia salina* metanauplii, and (iii) SuperFed (superfood diet – where corals were fed with enriched *Artemia salina* metanauplii). Measurements were performed after a two-week recovery period (Experiment 2- Phase 3), during which all corals were maintained at 26 °C. Temperature labels refer to the prior thermal exposure during Phase 2 (control: 26 °C; thermal stress: 32 °C). Data are presented as box plots showing the median, interquartile range, and individual data points. Different letters indicate significant differences (p ≤ 0.05) among dietary treatments at the same temperature history, while asterisks indicate significant differences (p ≤ 0.05) between temperature histories within the same dietary treatment.

#### 2.2) Diet modulates physiological traits and oxidative metabolism responses to thermal stress

At 32°C, only Non-Fed corals exhibited declines in chlorophyll content per surface area and per symbiont cell (Tukey, p < 0.018, Fig. 3b, c). Non-Fed corals also showed decreased symbiont density and the lowest ETRmax (Tukey, p < 0.03, Fig. 3a and e). Fed corals maintained symbiont density with the highest density among all diets (Tukey, p < 0.001, Fig. 3a), and also increased ETRmax (Tukey, p < 0.001, Fig. 3e). SuperFed corals also increased ETRmax (Tukey, p < 0.001, Fig. 3e) despite a decline in symbiont density (Tukey, p = 0.03, Fig. 3a). Superfed corals exhibited the highest ETRmax values among treatments after thermal stress (Tukey, p < 0.001, Fig. 3e). Protein content was unaffected by temperature or diet (Fig. 3d).

Oxidative stress biomarkers revealed distinct diet-dependent patterns under thermal stress conditions. Non-Fed corals were the only ones showing increased TAC (Tukey, p = 0.03, Fig. 4c) and LPO (Tukey, p < 0.0001, Fig. 4e), along with a reduced GSH/GSSG ratio (Tukey, p < 0.05, Fig. 4b). Fed corals exhibited elevated CAT activity (Tukey, p < 0.013, Fig. 4a) and reduced GSH/GSSG ratio (Tukey, p =0.008, Fig. 4 b) but no changes in LPO or TAC. SuperFed corals maintained control levels of LPO and TAC, but also preserved their GSH/GSSG ratio. Notably, they exhibited the highest antioxidant enzyme activity (CAT) and redox balance (GSH/GSSG) among all groups at 32 °C (Tukey, p < 0.001, Fig.4a and b).

#### 2.3) Enhanced recovery in Superfed corals

Following thermal stress, SuperFed corals exhibited enhanced recovery responses, particularly in photophysiological performance and oxidative status. While all heat-stressed corals exhibited reduced symbiont densities relative to corals maintained at 26°C (Tukey, p < 0.001, Fig. 5a), SuperFed corals maintained enhanced photophysiological performance, characterized by compensatory increases in chlorophyll content per skeletal surface area and per symbiont cell, higher ETRmax than all other diet treatments (Tukey, p < 0.001, Fig. 5b, c, f), and the lowest ROS levels (Tukey, p = 0.03, Fig.5e). Fed and Non-Fed corals exhibited increased chlorophyll per symbiont cell (Tukey, p < 0.001, Fig. 5c). Still, ETRmax was significantly reduced in Non-Fed corals (Tukey, p < 0.0001, Fig. 5f) while Fed corals showed decreased protein content compared to their control and SuperFed corals (Tukey, p < 0.02, Fig. 5d). ROS levels remained unchanged in Fed corals but decreased in Non-Fed corals (Tukey, p = 0.02, Fig.5e).

Two-way ANOVA statistics are detailed in Supplementary Tables S2 (Phase 2 – thermal stress) and S3 (Phase 3 – recovery). Summary schematics of physiological outcomes are presented in Figures 6 and 7.

**Figure 6.**
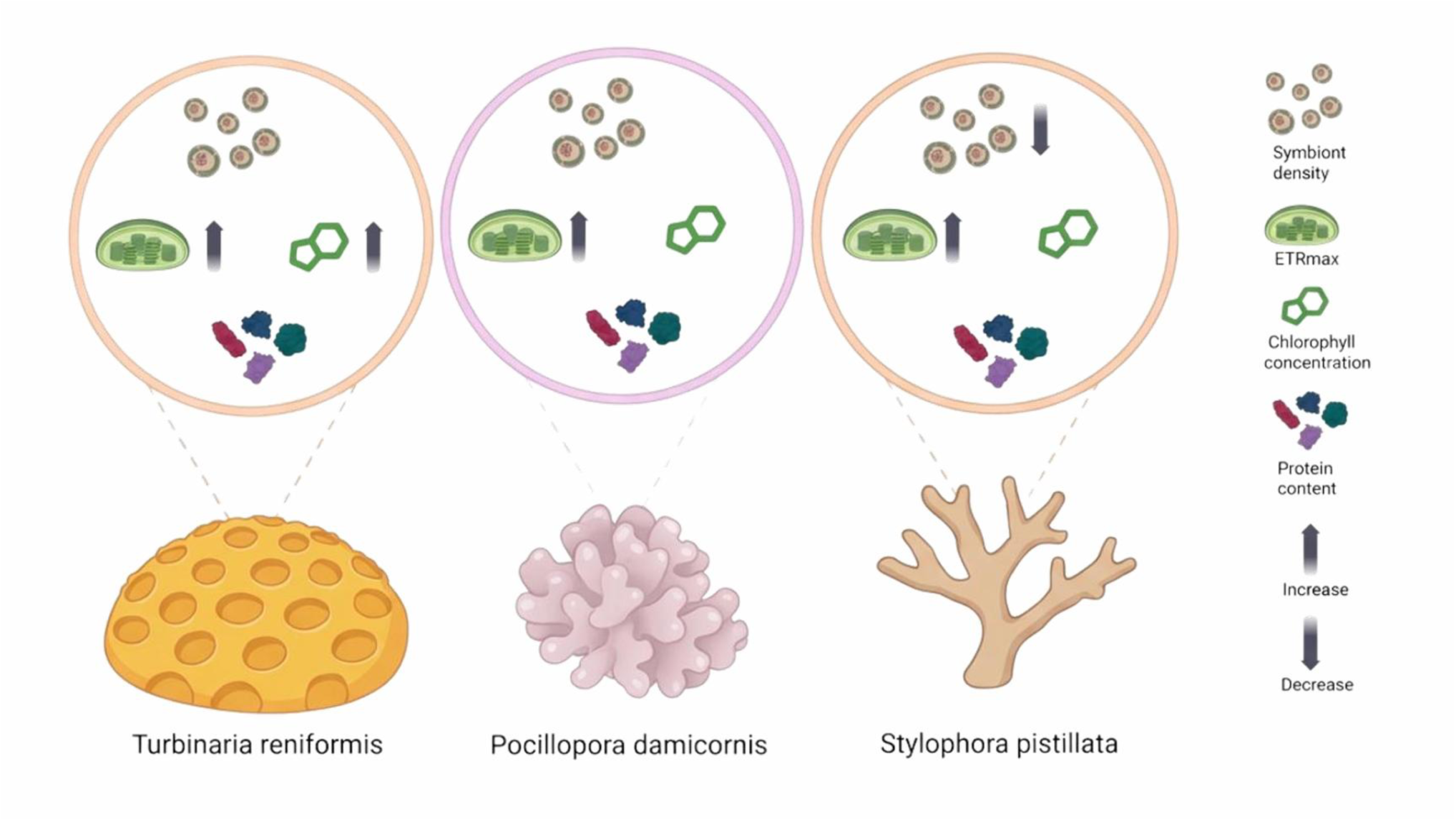
Overview of the physiological changes associated with antioxidant supplementation under thermal stress across three Red Sea coral species with different morphologies: *Turbinaria reniformis*, *Pocillopora damicornis*, and *Stylophora pistillata*. Measured metrics include symbiont density, chlorophyll concentration (chl *a* + *c2*), host protein content, and photophysiological performance (maximum electron transport rate, ETRmax). In *T. reniformis*, corals receiving antioxidant diet supplementation increased their chlorophyll concentration, and exhibited a 30% increase in ETRmax - over 55% higher efficiency than corals on the other treatments. In *P. damicornis,* supplemented corals maintained symbiont density and showed a 25% higher ETRmax than those on other diets. In *S. pistillata*, despite reductions in symbiont density, chlorophyll concentration did not decrease; supplemented corals showed a 97% increase in ETRmax compared to controls and were up to 70% higher than other diet groups. Arrows indicate an increase (↑) or decrease (↓), and the absence of arrows indicates no alteration.

**Figure 7.**
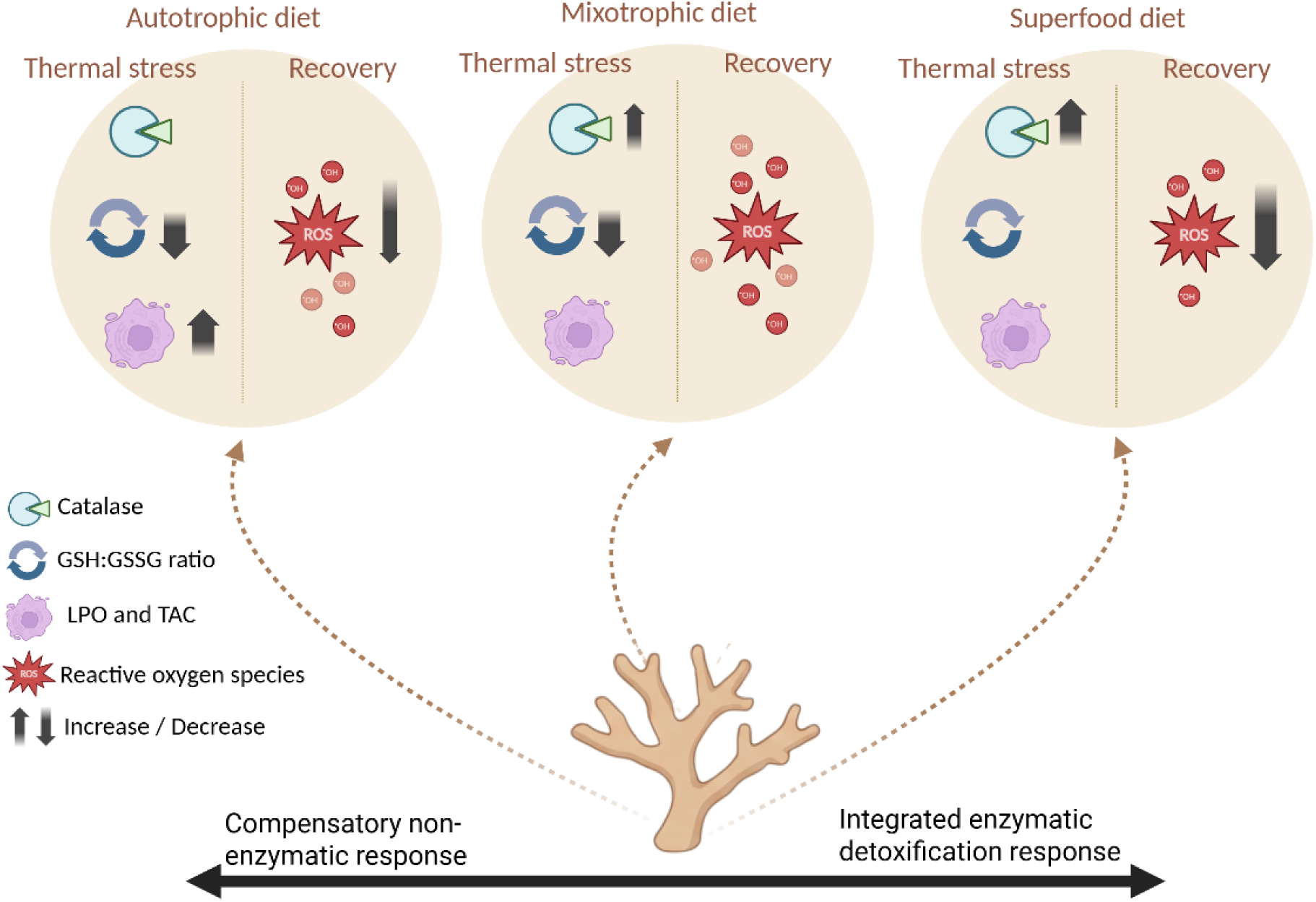
Summary of oxidative stress responses in *Stylophora pistillata* under thermal stress and recovery across different dietary treatments. Biomarkers include catalase activity (Catalase), reduced-to-oxidized glutathione ratio (GSH:GSSG), total antioxidant capacity (TAC), lipid peroxidation (LPO), and reactive oxygen species (ROS) levels. Arrows indicate increase (↑) or decrease (↓). Arrow thickness indicates response intensity. Absence of arrows indicates no alteration. Thermal stress induced oxidative stress in all diets, but with distinct profiles. Corals under the autotrophic diet showed the strongest stress signature, with a marked decline in GSH:GSSG, increased TAC response, and a 125% increase in lipid peroxidation. Corals in the mixotrophic diet-maintained lipid integrity by upregulating Catalase activity but still exhibited GSH:GSSG depletion and the highest ROS levels during recovery. In contrast, corals under antioxidant-enriched diet (Superfood diet) exhibited enhanced antioxidant capacity, with a ∼ 50% increase in catalase activity, stable GSH:GSSG ratios, no LPO increase, and ∼ 40% lower ROS levels during recovery compared to other diets.

## Discussion

In the face of escalating environmental pressures on coral reefs ^7^, our study demonstrates the potential of targeted antioxidant supplementation to enhance coral resilience. By enriching *Artemia salina* with antioxidants and other elements important for antioxidant metabolism, we developed a functional diet that improved photophysiological performance under thermal stress across three coral species and was further supported by enhanced antioxidant responses in *S. pistillata*. These findings highlight the value of enhancing coral nutrition—specifically through antioxidant support—as a strategy to reduce bleaching susceptibility under thermal stress conditions.

### The effect of diets under control conditions

The effects of heterotrophy on tissue parameters and photosynthetic processes varied among species under non-stressful conditions (26 °C). This agrees with the fact that reliance on heterotrophy and its physiological benefits can vary greatly among species ^41,42^ and with the amount of food available ^43,44,45^. In this study, an increased photophysiological performance (ETRmax) was observed only in the SuperFed corals of the species *T. reniformis*, and in both the Fed and SuperFed colonies of *P. damicornis* compared to the Non-Fed corals. In both species, this increase occurred without changes chl concentration, whereas in *P. damicornis* it was associated with lower symbiont density in SuperFed corals, suggesting a smaller but more efficient/healthier symbiont population. Heterotrophic feeding provides corals with essential nutrients such as nitrogen and phosphorus, alleviating nutrient limitation in their symbiotic algae and supporting more efficient photosynthetic function. By better meeting metabolic demands, it supports the repair and turnover of photosynthetic components, enabling enhanced electron transport and overall photosynthetic performance. Interestingly, *S. pistillata* under Fed and SuperFed diets showed slightly lower chlorophyll content than the Non-Fed corals, but no alteration in their photophysiological performance. Although carbon budget and photosynthetic rates were not directly assessed, this response may indicate a shift from photoautotrophy toward increased reliance on heterotrophy under higher zooplankton availability. Consistent with this interpretation, previous work has shown that higher feeding rates of *S. pistillata* were associated with a reduced autotrophic capacity ^47^. Neither mixotrophic nor superfood diets resulted in increased protein content or symbiont density in any of the three coral species, as is often reported in fed scleractinian corals ^48^. This may reflect the relatively short duration of the feeding treatment or a shift in energy allocation toward processes such as calcification rather than tissue growth ^47^. In addition, fed corals can also produce photosynthates of higher quality and energy yield ^48,49^.

Regarding oxidative stress, no differences were observed among diets in *S. pistillata* under control conditions for CAT activity, GSH/GSSG ratio, TAC, or LPO. This is consistent with similar ROS levels across treatments at 26 °C, ruling out metabolic disorders related to oxidative imbalance due to unhealthy dietary treatments ^50^.

### Responses to thermal stress under different diets

The superfood diet was associated with improved photophysiological performance under thermal stress across the coral species studied, as reflected by higher ETRmax values compared to the other diet treatments. The antioxidant-enriched diet likely limited the accumulation of ROS generated under heat-stressed photosynthesis, thereby reducing oxidative stress pressure on the photosynthetic apparatus. By preventing excessive ROS buildup, antioxidants help preserve the integrity of photosystem II components while simultaneously supporting the repair and turnover of damaged proteins and pigments. This balance between damage and repair allows electron transport to occur more efficiently, preventing photoinhibition and sustaining higher photosynthetic performance ^51^, which is reflected as a sustained high ETRmax even under thermal stress.

More specifically, the enhancement of photophysiological performance in SuperFed corals may be partly attributed to the presence of manganese and ascorbate (vitamin C) in the diet. Previous studies have shown that manganese acts as an antioxidant and can increase the photosynthetic efficiency and overall photosynthetic rate of corals under both control and heat stress conditions ^22,52^. In turn, ascorbate plays a crucial role in photosynthesis, participating in key biochemical processes, including: (i) scavenging H_2_O_2_ produced during photoreduction in PSI via the action of ascorbate peroxidase (AP), and (ii) regulating photosynthetic electron transport through the monodehydroascorbate (MDA) radical, which is formed by AP and can serve as a direct electron acceptor for PSI (for a review, see ^53^). These mechanisms provide a plausible explanation for the observed responses, although they were not directly assessed in this study.

Interestingly, thermally stressed *T. reniformis* corals fed the superfood diet exhibited the highest ETRmax across treatments, while no reduction in symbiont density was observed in any treatment group. Although Fed corals showed an early bleaching signature through reduced chlorophyll content per skeletal surface area, SuperFed corals showed increased chlorophyll content per symbiont cell under thermal stress. Together, these responses suggest a potential optimization of the photosynthetic apparatus and enhanced photophysiological resilience before the onset of bleaching in this species.

*Pocillopora damicornis* corals fed the superfood diet also exhibited the highest ETRmax across treatments and maintained symbiont densities comparable to control conditions. Superfood may have helped maintain cellular homeostasis and reduce stress signaling that typically leads to symbiont expulsion and lower photophysiological performance^10,14^. By mitigating oxidative stress and sustaining metabolic demand, the host–symbiont partnership remained stable, allowing symbiont densities to be maintained. In addition, the combination of (i) lower symbiont density in SuperFed corals before thermal stress, and (ii) higher photophysiological performance under thermal stress, may have reflected a more efficient symbiont assemblage, potentially reducing the risk of excess oxidative pressure. Consistent with this interpretation, previous studies have shown that reduced symbiont densities limit the accumulation of ROS under stress conditions ^55^. In contrast, the corals under other dietary treatments exhibited rather reductions in symbiont density or lower photophysiological performance during thermal stress, indicative of stress response.

Finally, both Fed and SuperFed colonies of *S. pistillata* showed enhanced photosynthetic performance under thermal stress. Specifically, ETRmax increased by 25% in Fed corals and by an impressive 75% in SuperFed corals compared to Non-Fed corals. These observations align with previous studies, which showed that heterotrophy can increase symbiont density and photosynthetic efficiency in *S. pistillata* under heat stress ^46^. The additional increase observed in SuperFed corals suggests a further improvement in symbiont photophysiological function as described above for the other coral species.

### Diet-dependent shifts in antioxidant defense strategies under thermal stress and recovery

In this study, oxidative stress biomarkers were assessed in *S. pistillata* to provide biochemical context for the physiological responses observed across species. Concerning the enzymatic antioxidant system, CAT has a high *K_m_* value for H₂O₂, making it particularly effective at scavenging high concentrations of ROS ^25^. In fact, CAT has the highest turnover rate among all antioxidant enzymes, decomposing H₂O₂ rapidly and efficiently ^56^. Its activity has been linked to coral bleaching ^57,58^. In *S. pistillata*, CAT activity increased under thermal stress, with SuperFed corals exhibiting CAT activity approximately 50% greater than Non-Fed and 40% greater than Fed corals. Notably, thermal stress induced a 150% increase in CAT activity in SuperFed corals compared to control corals at 26 °C, whereas increases did not exceed 100% in the other diet treatments. These results indicate an enhanced capacity of SuperFed corals to upregulate CAT activity under thermal stress, a key mechanism involved in ROS detoxification and cellular protection ^59^. The enhancement of CAT activity in SuperFed corals may be linked to the higher availability of micronutrients such as iron in their diet, an essential component of CAT ^60^. Indeed, previous studies have reported increased cellular demand for iron to sustain high CAT activity ^61^ and correlations between iron availability and CAT activity ^62^.

Regarding the non-enzymatic antioxidant system, corals use glutathione (GSH) to help maintain a reducing environment within cells, which is essential for protection against oxidative stress ^25^. Glutathione acts as an antioxidant by reacting with ROS like singlet oxygen (¹O₂), anion superoxide (O₂⁻), and hydroxyl radicals (HO•) and changing from its reduced (GSH) to its oxidized form (GSSH) ^63^. Consequently, the GSH/GSSG ratio is widely used as a marker of oxidative stress in cells, with decreases indicating oxidative stress. SuperFed corals were the only ones that maintained a stable GSH/GSSG ratio, while corals on the other diets showed increased oxidation of the glutathione pool.

Decreased photosynthetic efficiency has been reported in thermally sensitive Symbiodiniaceae species, such as *Breviolum minutum* and *Effrenium voratum* ^64^, to be related to increased glutathione oxidation ^59^. Consistently, our results suggest that the GSH-enriched *Artemia salina* metanauplii may have contributed to maintaining glutathione redox homeostasis, potentially by supporting intracellular GSH availability ^65^.

Overall, SuperFed corals showed both enhanced CAT activity and maintenance of glutathione redox balance under thermal stress, which was associated with higher photophysiological performance. These findings are consistent with previous research highlighting the importance of antioxidant regulation in mitigating oxidative damage and sustaining photosynthetic function under stress ^66^. In addition, glutathione redox homeostasis has been associated with improved thermal tolerance and photosynthetic efficiency in Symbiodiniaceae ^59^. Neither Fed nor SuperFed corals exhibited an increase in non-enzymatic total antioxidant capacity (TAC), yet no increase in lipid peroxidation was observed. This suggests that enzymatic antioxidant responses may have been sufficient to limit oxidative damage, potentially reducing the need for mobilization of non-enzymatic antioxidant reserves. In particular, the strong upregulation of CAT activity in Fed (∼ 2-fold) and SuperFed (∼ 2.5-fold) corals may have contributed to limiting lipid damage under thermal stress.

In contrast, Non-Fed *S. pistillata* corals were the only treatment to upregulate TAC under thermal stress, along with a smaller, non-significant increase in CAT activity. However, these responses were accompanied by an approximately 50% rise in lipid peroxidation in their symbionts, indicating the occurrence of oxidative damage. Together, these findings suggest that corals under a predominant autotrophic diet adopted a distinct antioxidant strategy characterized by increased reliance on non-enzymatic antioxidant defenses, potentially reflecting the higher metabolic costs associated with sustained enzymatic antioxidant activity^25,27^. Nevertheless, this response was insufficient to mitigate oxidative damage to lipids under thermal stress. This vulnerability may also be related to differences in lipid composition. Previous studies showed that polyunsaturated fatty acids (PUFAs) can enhance membrane thermal stability and reduce susceptibility to reactive oxygen species (ROS) ^67, 32^. As these lipids are commonly obtained through heterotrophic feeding, their reduced availability in non-fed corals may have contributed to the observed increase in oxidative damage.

Overall, we observed a clear gradient in antioxidant strategy, ranging from greater reliance on non-enzymatic defenses in Non-Fed corals, marked by increased TAC and glutathione pool oxidation, to an intermediate response in Fed corals, characterized by increased CAT activity but reduced GSH:GSSG ratio, and finally to a more efficient enzymatic antioxidant response in SuperFed corals, which maintained glutathione redox homeostasis. Given the substantial energetic and micronutrient demands associated with antioxidant enzyme synthesis and activity, this pattern suggests that enhanced nutritional quality may shift antioxidant regulation from a compensatory non-enzymatic response toward a more efficient and integrated enzymatic detoxification strategy under thermal stress.

After the 2-week recovery period, *S. pistillata* corals appeared to compensate for the lower symbiont density following thermal stress by increasing chlorophyll concentration per symbiont cell, however, only SuperFed ones were able to increase the chlorophyll concentration per skeletal surface area. Maintaining a smaller symbiont population during the recovery phase may be advantageous, as previously suggested ^55^. Notably, SuperFed corals had the lowest ROS levels – 45% lower than corals under the other diets. This reduction in ROS was accompanied by higher photophysiological performance, with ETRmax values up to 60% higher than in corals under other diets and 50% higher than the corresponding control, suggesting that improved regulation of ROS levels may be linked to enhanced photosynthetic function ^66^. Notably, this recovery pattern mirrored the antioxidant responses observed during thermal stress, with Non-Fed corals showing impaired photophysiological recovery, Fed corals maintaining intermediate performance, and SuperFed corals exhibiting enhanced recovery above control levels.

Corals under a strictly autotrophic diet (Non-Fed) were the only treatment exhibiting reduced photophysiological performance during recovery, supporting the idea that heterotrophic supplementation can enhance photophysiological performance during recovery, potentially through positive feedback between heterotrophic and autotrophic processes ^68^. Fed corals showed depleted protein content at the end of the recovery period, suggesting depletion of host reserves. A similar pattern has been observed for *Porites compressa*, which depleted energy reserves during bleaching recovery despite an increase in chlorophyll and photosynthesis and the presence of zooplankton ^69^. Our results further suggest that coral species can adjust their recovery responses depending on the quality and availability of food resources, with SuperFed corals maintaining both protein content and higher photophysiological performance compared to corals under a non-enriched diet.

In sum, the dietary treatments revealed a clear gradient in antioxidant regulation, ranging from insufficient antioxidant protection in Non-Fed corals, to partial protection in Fed corals characterized by increased CAT activity but persistent redox imbalance, and finally to enhanced oxidative homeostasis in SuperFed corals, which combined strong CAT upregulation with stable GSH:GSSG ratios and reduced ROS accumulation during recovery.

### Constraints and perspectives on coral resilience

Our findings highlight the potential of antioxidant-enriched feeding to support coral physiological performance under thermal stress, particularly through modulation of oxidative stress responses. These results suggest that targeted enhancement of antioxidant capacity may represent a promising strategy to improve coral condition during periods of environmental stress. However, some limitations should be considered. Biochemical assessments of oxidative stress were conducted only in *S. pistillata*, providing a mechanistic insight for this species while limiting direct extrapolation across taxa. In addition, the study was performed under controlled laboratory conditions, and further research is needed to determine whether these responses persist under more complex and variable conditions of natural reef environments.

From an applied perspective, targeted nutritional supplementation may offer a practical approach to support coral performance in controlled settings, particularly in *ex situ* (land-based) nurseries where feeding regimes and environmental conditions can be regulated. Implementation *in situ* nurseries may be more challenging due to logistical constraints and environmental variability. Nevertheless, emerging technologies may help overcome some of these limitations by enabling targeted delivery of nutritional supplements in field settings, as discussed in a recent perspective on the role of nutrition in coral restoration scalability ^36^.

Future research should incorporate time-resolved measurements to better characterize acclimation and recovery dynamics, while also investigating species-specific nutritional requirements, optimization of nutrient formulations, and potential interactions with complementary intervention strategies such as microbiome manipulation and selective breeding. Assessing delivery methods across life stages and across both *ex situ* and *in situ* contexts will be critical for translating these findings into scalable restoration practices.

## Materials and Methods

### Antioxidant-enriched food preparation (Superfood)

Superfood was produced by incubating *Artemia salina* metanauplii in filtered seawater enriched with antioxidants, essential metals, and lipids (details on the supplements and concentrations selected are in table 1). First, the *Artemia salina* cysts were hatched according to the manufacturer instructions (TQ Artemia, INVE Aquaculture®, Belgium) and transferred to a solution (seawater with dissolved supplements) for an additional 12h incubation. The incubations were carried out under constant aeration to avoid mortality. After enrichment, *Artemia salina* metanauplii (∼ 400 µm) were harvested, counted, and then portioned into aliquots, which were subsequently frozen for later use. Non-enriched *Artemia salina* metanauplii (Fed treatment) were subjected to the same hatching, incubation, and handling procedures as the enriched group, but without the addition of antioxidant supplements during the enrichment phase. At each feeding event, approximately 1000 *Artemia salina* metanauplii were provided per coral nubbin. During the feeding, the inflow of seawater was stopped for one hour to allow prey capture, while the submersible pumps remained on to maintain water circulation and prevent settling of prey. After the 1 h feeding period, the flow-through system was resumed, ensuring rapid water renewal and removal of excess food.

**Table 1.** Supplements and concentrations used in our targeted antioxidant supplementation and their known/potential benefits for symbiotic corals.

| Supplement | Concentration | Known/Potential Benefits | References |
| --- | --- | --- | --- |
| Retinol (Vitamin A) | 2.75 mg/L | Supports cell growth, differentiation, tissue repair, and immune response. | 70, 71, S. Presso Selco INVE Aquaculture. |
| Sodium Ascorbate (Vitamin C) | 8 mg/L | Antioxidant protecting against oxidative stress; regulates photosynthetic electron transport. | 53, S. Presso Selco INVE Aquaculture. |
| α-Tocopherol (Vitamin E) | 0.13 mg/L | Lipid-soluble antioxidant protecting against lipid peroxidation, promotes membrane stability. | 72, 73, S. Presso INVE Selco. |
| L-Glutathione (Reduced) | 1.84 g/L | Non-enzymatic antioxidant regulating redox homeostasis; enhances cellular immunity and feeding behavior. | 59, 74. |
| Astaxanthin | 4 mg/L | UV protection; reduces photodamage; enhances pigmentation and may improve photosynthesis in symbiotic algae. | 75, 76, Algamac Enrich Aquafauna Biomarine Inc. |
| DHA (Docosahexaenoic Acid) | 86 mg/L | Omega-3 fatty acids support membrane fluidity and function. | 67, Algamac 350 Aquafauna Biomarine Inc. |
| EPA (Eicosapentaenoic Acid Methyl Ester) | 5.8 mg/L | Omega-3 fatty acids support immune function and growth; maintain membranes and reduce inflammation. | 77, Algamac 350 Aquafauna Biomarine Inc. |
| Manganese (Mn, as Manganese Chloride) | 0.4 μg/L | Cofactor in enzymes for photosynthesis and oxidative stress protection; supports nutrient cycling. | 22, 52. |
| Magnesium (Mg, as Magnesium Chloride) | 0.2 g/L | Stabilizes aragonite structure; involved in pH regulation of the coral internal medium. | 78. |
| Iron (Fe, as Iron III Chloride) | 1.5 μg/L | Essential for photosynthesis; participates in energy production and antioxidant enzyme activity. | 22, 79. |

### 1. Experimental design

The experiments were performed with corals originally collected in the Gulf of Aqaba, Red Sea, under the Convention on International Trade in Endangered Species of Wild Fauna and Flora license number DCI-89-32 and maintained in closed aquaria at the Centre Scientifique de Monaco (CSM). Species were selected to represent ecologically relevant coral taxa with contrasting morphologies and physiological responses to environmental stress. *Turbinaria reniformis* and *Pocillopora damicornis* were used to assess treatment effects across species with differing structural and functional traits, allowing broader evaluation of responses. *Stylophora pistillata*, a widely used model species in coral experimental research, was selected for more detailed physiological analyses, including recovery dynamics and oxidative stress responses.

Six different colonies of *T. reniformis* and *P. damicornis* were used to generate 6 nubbins per colony, while 6 colonies of *S. pistillata* were used to generate 24 nubbins per colony. Coral nubbins (4-6 cm in length) were randomly distributed and kept into 12 independent 20 L experimental tanks supplied with natural seawater (flow rate of 10 L h^-1^).

Metal halide lamps (Philips, HPIT 400 W, Distrilamp®, France) provided irradiance of 250 µmol photons m^-2^ s^−1^ (photoperiod of 12:12 light:dark). The seawater temperature was maintained at 26 ± 0.5 °C using submersible heaters (Visi-TermH Deluxe, Aquarium Systems®, France) and continuously monitored using an automated control and logging system to ensure stable thermal conditions. Salinity was constant at 38 PSU. Submersible pumps ensured thorough mixing of the water, and the aquaria were cleaned regularly on a weekly basis to prevent algal overgrowth. Continuous flow-through conditions (10 L h^-1^) ensured regular water renewal and stable water quality throughout the experiment.

Coral nubbins were acclimated to the experimental tanks for 3 weeks under standard husbandry conditions used at the CSM, including regular feeding with *Artemia salina* metanauplii at repletion twice a week to maintain coral health and physiological stability prior to the experimental treatments.

After acclimatization, two thermal stress experiments were carried out as described below. Thermal stress levels were adjusted according to species-specific tolerance, with higher temperatures applied to *S. pistillata* to ensure a comparable level of physiological challenge across species.

*Experiment 1* - *T. reniformis* and *P. damicornis* were subjected to three dietary treatments (Phase 1 – Feeding, with four replicate tanks per condition): (1) Autotrophic diet (Non-Fed), in which the corals relied solely on light and nutrients naturally present in seawater, without any additional food supplementation, in an effort to replicate a highly oligotrophic environment where the coral nutritional needs are largely met by their symbionts; (2) Mixotrophic diet (Fed), in which the corals were fed non-enriched *Artemia salina* metanauplii 3 times a week, to simulate a less oligotrophic reef environment where corals also have a significant nutritional contribution from heterotrophic feeding on plankton; and (3) Superfood diet (SuperFed), in which corals were fed superfood (enriched *Artemia salina* metanauplii) 3 times a week. After 2 weeks under these conditions, half of the tanks from each dietary treatment were subjected to thermal stress, resulting in a total of six treatments (Phase 2 - Thermal stress, with two replicate tanks per condition). The temperature was raised from 26 ± 0.5 °C to 30 °C over a period of 5 days (0.8 °C per day, following recommendations in ^80^) and maintained at 30 ± 0.5 °C for a further 7 days. No feeding occurred during the thermal stress phase. At the end of Phase 2, coral nubbins (N = 6 per species, 3 per tank) were collected and evaluated for photophysiological performance (maximum electron transport rate, ETRmax) and bleaching descriptors (chlorophyll a and c_2_ concentration, host protein content, and symbiont density). Graphical details on Experiment 1 are available in the supplementary material (Figure S1).

*Experiment 2* - *Stylophora pistillata* was subjected to the same dietary treatments as described in Experiment 1. Nubbins were randomly distributed into 12 aquaria, with three nubbins per colony per tank. After 2 weeks of feeding (Phase 1), thermal stress was applied to half of the tanks of each dietary treatment (Phase 2). Due to the higher thermal tolerance of *S. pistillata*, the temperature was increased from 26 ± 0.5 °C to 32 °C over a period of 7 days (0.8 - 0.9 °C per day) and maintained at 32 ± 0.5 °C for an additional 2 weeks. No feeding occurred during the thermal stress phase. Following thermal stress, the temperature gradually lowered to 26 °C, and the feeding treatments were resumed for two more weeks to assess the recovery of the corals (Phase 3 - Recovery). At the end of phases 2 and 3, nubbins (N = 6) were used to measure the photophysiological performance and bleaching descriptors, as described above. Enzymatic and non-enzymatic oxidative stress biomarkers (N = 6) were assessed after Phase 2, and ROS levels were quantified after the recovery phase (Phase 3). Graphical details on Experiment 2 are available in the supplementary material (Figure S2).

### 2. Physiological analyses

#### 2.1. Bleaching descriptors: chlorophyll a and c_2_ concentration, symbiont density, and protein content

Coral nubbins of the three coral species (N = 6 per treatment for each species) were frozen at −80 °C at the end of Phase 2 (thermal stress) of each experiment to determine chlorophyll (*a* and c_2_), symbiont density, and coral host protein content according to ^81^. Tissue was removed from the skeleton and collected in 10 mL of 0.45 µM FSW using an airbrush. The resulting tissue slurry was homogenized with a potter grinder. Sub-samples were used to quantify symbiont density on a Z1 Coulter Counter (Beckman Coulter) and to determine the protein content of the coral host using the Bradford Protein Kit (23200, Thermo Fisher Scientific, USA). The chlorophyll (chl) concentration was determined according to ^82^. For this purpose, 5 mL of the homogenized tissue slurry was centrifuged (8,000 *g*, 10 min), the supernatant was removed, and the symbionts were resuspended in 5 mL of acetone for the extraction of chlorophyll *a* and c_2_ under dark, cold conditions for 24h. Data were normalized to surface area (cm^2^) using the wax-dipping method ^83^.

#### 2.2. Fluorometry

A Pulse Amplitude Modulation fluorometer (Walz®, Germany) was used to determine the minimal (F0) and maximal (Fm) chlorophyll a fluorescence of each nubbin acclimated to the experimental conditions (n=6 nubbins per treatment and species). After a dark adaptation of 20 min, the initial fluorescence level (F_0_) was determined by applying weak modulated pulses of red measuring light (MI=10). A 0.8 s saturating pulse of actinic light (SI=10 corresponding to an intensity of 4000 µmol photons m^-2^ s^-1^) was then applied to measure the maximum fluorescence level (F_m_). Dark-adapted PSII rapid-light curves (RLC) were then measured for each colony according to ^84^ at irradiances of 0, 11, 18, 27, 42, 75, 131, 221, 665, 1033, and 1957 mmol photons m^-2^ s^-1^. PSII parameters obtained during the RLC, such as the relative electron transport rate (rETR) were used to study the PSII activity. ETRₘₐₓ is widely considered a reliable proxy for coral photosynthesis and photosynthetic efficiency, as it reflects the maximum capacity of the photosynthetic electron transport chain and correlates strongly with carbon fixation rates under saturating light conditions. Due to the time required to process all samples, measurements were conducted sequentially within a consistent daily time window (12:00 – 15:00 h). Replicates from each treatment were measured in alternating sequence to ensure balanced comparison across treatments.

### 3. Oxidative metabolism analysis in Stylophora pistillata

#### 3.1. Sample preparation

To perform the biochemical assays, 6 nubbins of *S. pistillata* from each experimental treatment were snap-frozen in liquid nitrogen immediately after collection and subsequently stored at −80 °C. Samples were prepared as described in ^85^. Briefly, coral subsamples were cut from each nubbin (∼ 0.5 cm^2^) and homogenized in ice with sonication (Frequency 70 kHz, Vibra-Cell™ 75,185, Bioblock Scientific®, France) using 250 – 300 µL of the specific homogenization buffer for each analysis, as described below. After sonication, the remaining skeleton was discarded, the homogenized holobiont solution was centrifuged, and the intermediary phase was collected and immediately used for analysis. The total protein content of holobiont homogenates was determined according to ^86^ using the “Comassie (Bradford) Protein Assay Kit” (Thermo Scientific®, USA).

#### 3.2. Catalase activity

The measurement of catalase (CAT) activity was performed in *S. pistillata* after Phase 2 (thermal stress) of the experiment according to the Catalase Activity Assay (ab83464, Abcam®, United Kingdom) protocol. The catalase present in the samples reacts with H_2_O_2_ to produce water and oxygen. The unconverted H_2_O_2_ reacts with a probe to generate a product measured calorimetrically at 570 nm using a microplate reader (Xenius®, Monaco) in a 96-well flat-bottom microplate. Catalase activity present in the sample is inversely proportional to the signal obtained. Data were normalized to the total protein content in each well and expressed as mU (nmol min^−1^) mg^−1^ protein.

#### 3.3. Glutathione concentration

Reduced (GSH) and oxidized (GSSG) glutathione concentrations were determined in *S. pistillata* after Phase 2 (thermal stress) of the experiment using a commercial reagent kit (38185, Sigma-Aldrich®, USA) following the manufacturer’s instructions. Absorbance readings at 415 nm were conducted in a 96-well flat-bottom transparent microplate using spectrofluorometry (Xenius®, Monaco). Data were normalized to the total protein content in each well and expressed as mmol mg^-1^ protein, and the GSH/GSSG ratio, which provides a reliable estimation of cellular redox status, was calculated.

#### 3.4. Non-enzymatic Total Antioxidant Capacity

The determination of non-enzymatic total antioxidant capacity (TAC) of soluble low molecular weight antioxidants was conducted in *S. pistillata* after experiment Phase 2 (thermal stress) using the OxiSelect® Total Antioxidant Capacity (TAC) Assay Kit, following the manufacturer’s instructions. The antioxidant net absorbance values of the samples were compared with a known uric acid standard curve, with absorbance values being proportional to the total antioxidant capacity of the sample. Absorbance readings were conducted in a 96-well flat-bottom transparent microplate using spectrofluorometry (Xenius®, Monaco). Data were normalized to the total protein content in each well and expressed as μmol L^−1^ copper reducing equivalents (CRE) mg^−1^ protein.

#### 3.5. Lipid Peroxidation

The oxidative damage to lipids (lipid peroxidation, LPO) was determined in *S. pistillata* after Phase 2 (thermal stress) according to the method described by ^87^. Measurements were conducted in the coral host and symbiont fractions separately. For that, the coral host fraction was separated from symbionts following the procedure described by ^55^. Samples were homogenized in KCl (1.15%) solution containing 35 µmol L^−1^ butylated hydroxytoluene (BHT), and centrifuged for 10 min (10,000 *g*, 4 °C). Samples (20 µL) were added to a reaction mixture containing BHT solution (35 µmol L^−1^), 20% acetic acid (pH 3.5), TBA (0.8%), sodium dodecyl sulfate (8.1%), and ultrapure water, and then incubated in a water bath at 95 °C for 30 min. After cooling, n-butanol was added with thorough vortexing. Mixtures were centrifuged (300 rpm, 15 °C, 10 min), and the immiscible organic layer was collected and added to a 96-well flat-bottom black microplate. Fluorescence (excitation: 515 nm; emission: 553 nm) was measured using a spectrofluorometer (Xenius®, Monaco). Data were normalized to the total protein content in each well and expressed as nmol MDA mg^−1^ protein.

#### 3.6. Reactive oxygen species

The quantification of intracellular reactive oxygen species (ROS) was performed in *S. pistillata* at the end of Phase 3 (recovery). Measurements were conducted using the fluorescence technique outlined by ^88^, with modifications as described by ^85^. Each sample was homogenized in a buffer containing Tris-HCl (100 mmol L^−1^, pH 7.7), ethylenediaminetetraacetic acid (2 mmol L^−1^), and MgCl_2_ (5 mmol L^−1^), followed by centrifugation (20,000 *g*, 20 min, 4 °C). The protein content was adjusted to a final concentration of 0.8 mg mL^−1^ for both coral host and endosymbiont homogenates to optimize fluorescence curves over time. In a flat-bottom black microplate, 10 µL of sample was added to a medium containing HEPES (30 mmol L^−1^), KCl (200 mmol L^−1^), and MgCl_2_ (1 mmol L^−1^, pH 7.2). Subsequently, 10 µL of the fluorescent probe H_2_DCFDA (16 µmol L^−1^, Invitrogen) was added. Fluorescence (excitation: 488 nm; emission: 525 nm) was measured at intervals of 5 minutes for up to 50 minutes using a spectrofluorometer (Xenius®, Monaco). Results were expressed as fluorescence units per minute (F.U. min^−1^).

### 4. Statistical analysis

All physiological and biochemical data are presented as box plots with individual data points, showing median, interquartile range, and data distribution. Tank effects were tested by initially including tank as a factor in the model, according to ^89^. As no significant tank effect was detected, this factor was excluded, and tanks were pooled for subsequent analysis. For Phase 2 (thermal stress), the effects of feeding and temperature on physiological and biochemical parameters were evaluated using two-way analysis of variance (ANOVA), with temperature (two levels: 26 and 30°C for Experiment 1; 26 and 32°C for Experiment 2) and diet (three levels: Non-Fed, Fed, and SuperFed) as fixed factors. Data from Phase 3 (recovery) of Experiment 2 were also evaluated using two-way ANOVA, with temperature history (two levels: 26 °C, non-stressed; and 32 °C, previously heat-stressed) and diet (three levels: Non-Fed, Fed, and SuperFed) as fixed factors. Data residuals were visually inspected in R for normality and homoscedasticity using Q–Q plots versus fitted value plots. When significant effects were detected, Tukey’s post hoc tests were performed. Differences were considered statistically significant when p ≤ 0.05. All analyses were performed in R (R Core Team, 2023) within RStudio (version 2025.09.0+387). Figures were created using BioRender (BioRender.com).

## Supporting information

Marangoni et al Supplemental material

## Acknowledgments

A special thanks to Isabelle Nakaima for her invaluable assistance in creating the graphical figures.

## Funding

Centre Scientifique de Monaco (CSM, Monaco)

Murray Foundation (Mf, UK)

Coral Research & Development Accelerator Platform (CORDAP)

Coral Vivo Project (Programa Petrobras Socioambiental and Arraial d’Ajuda Eco Parque, Brazil).

## Author contributions

Conceptualization: L.F.B.M., C.F-P., M.M., O.L., M.L.

Methodology: L.F.B.M., E.B., M.C., M.M., M.L.

Validation: L.F.B.M., E.B., M.C., M.L.

Formal analysis: L.F.B.M.

Data curation: L.F.B.M.

Writing-original draft: L.F.B.M.

Writing-review & editing: L.F.B.M., C.F-P., M.M, O.L., M.L.

Visualization: L.F.B.M., C.F-P., M.M, O.L., M.L.

Supervision: C.F-P.

## Data and materials availability

The datasets analyzed during the current study are within the manuscript and its Supplementary Information files.

