## Supplementary material for "Improved coral thermal tolerance through modulation of antioxidant defenses": Marangoni et al Supplemental material

Laura F.B. Marangoni *et al.*

**This PDF file includes:**

Supplementary Text
Figs. S1 and S2
Tables S1 to S3

**Fig. S1.**

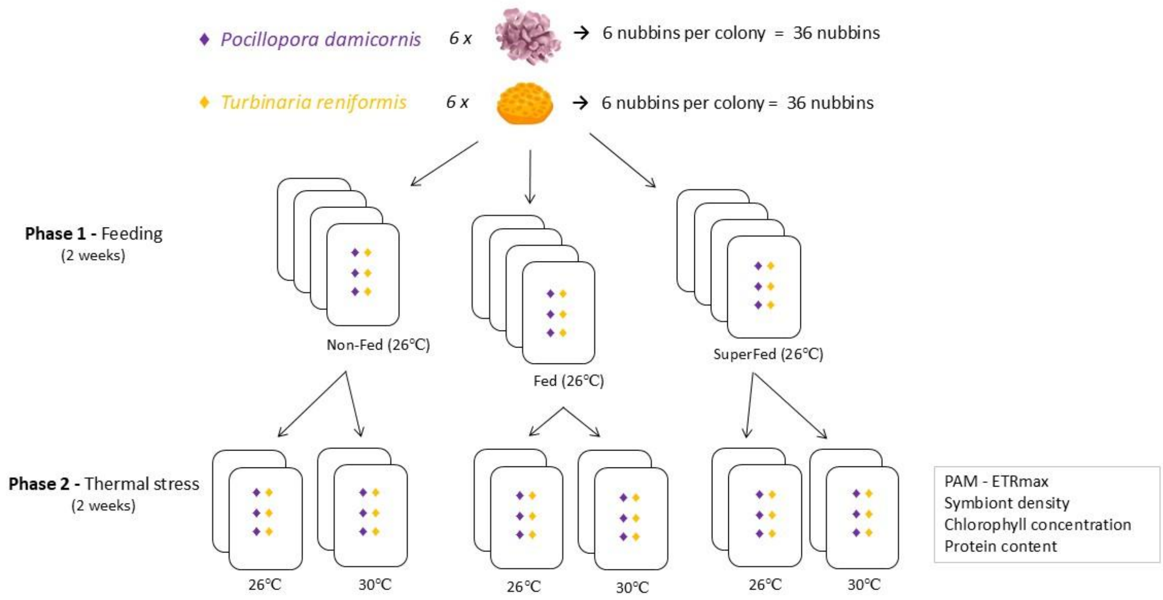


**Fig. S1. Experimental set-up for Experiment 1**: *Turbinaria reniformis and Pocillopora damicornis* nubbins were kept for 2 weeks under different diets (Phase 1 - Feeding) and then exposed to thermal stress for 2 weeks (Phase 2 – Thermal stress). Feeding treatments (diets) were as follows: (i) autotrophic diet - where corals were not fed (Non-Fed), (ii) mixotrophic diet - where corals were fed non-enriched *Artemia salina* metanauplii (Fed), and (iii) superfood diet – where corals were fed with enriched *Artemia salina* metanauplii (SuperFed). At the end of the experiment, nubbins (N = 6 per species, 3 per tank) were collected and evaluated for photosynthetic efficiency and bleaching descriptors (chlorophyll-*a* and $c_2$ concentration, host protein content, and symbiont density).

**Fig. S2.**

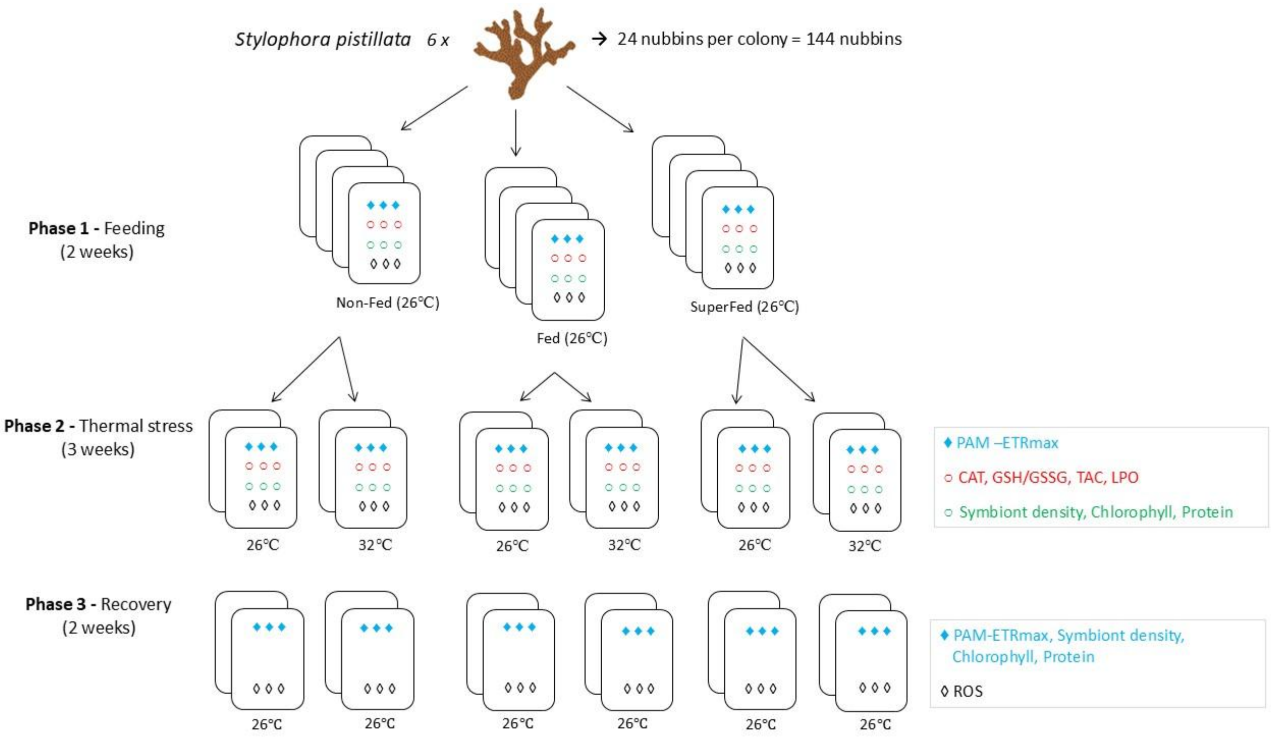


**Fig. S2. Experimental Setup for Experiment 2**: *Stylophora pistillata* nubbins were kept under different feeding treatments (diets) for 2 weeks (Phase 1 - Feeding), exposed to thermal stress for 3 weeks (Phase 2 – Thermal stress), and then submitted to a 2-week recovery period (Phase 3 - Recovery). Feeding treatments (diets) were as follows: (i) autotrophic diet - where corals were not fed (Non-Fed), (ii) mixotrophic diet - where corals were fed non-enriched *Artemia salina* metanauplii (Fed), and (iii) superfood diet – where corals were fed with enriched *Artemia salina* metanauplii (SuperFed). At the end of phases 2 and 3, nubbins (N = 6) were used to measure the photosynthetic efficiency and bleaching descriptors (chlorophyll-*a* and $c_2$ concentration, host protein content, and symbiont density). Oxidative stress biomarkers (N = 6) - Catalase (CAT) activity, reduced and oxidized glutathione ratio, non-enzymatic total antioxidant capacity (TAC), and lipid peroxidation (LPO) - were assessed after Phase 2. Reactive Oxygen Species (ROS) were quantified after the recovery period (Phase 3). Different shapes and color signs indicate the nubbins used for each analysis in each phase of the experiment.

**Table S1.**

| Parameter | Effect | *Pocillopora damicornis* | | | *Turbinaria reniformis* | | |
|---|---|---|---|---|---|---|---|
| | | ***df*** | ***F*** | ***p*** | ***df*** | ***F*** | ***p*** |
| Symbiont density | Temp | 1 | 9.42 | **0.005** | 1 | 15.9 | **<0.001** |
| | Diet | 2 | 1.82 | 0.178 | 2 | 0.09 | 0.917 |
| | Temp*Diet | 2 | 5.36 | **0.010** | 2 | 0.06 | 0.945 |
| Chlorophyll ($cm^2$) | Temp | 1 | 3.56 | 0.069 | 1 | 18.4 | **<0.001** |
| | Diet | 2 | 2.03 | 0.149 | 2 | 0.48 | 0.623 |
| | Temp*Diet | 2 | 1.35 | 0.274 | 2 | 4.26 | **0.023** |
| Chlorophyll (cell) | Temp | 1 | 8.35 | **0.010** | 1 | 3.33 | 0.078 |
| | Diet | 2 | 0.57 | 0.620 | 2 | 0.42 | 0.662 |
| | Temp*Diet | 2 | 3.63 | 0.060 | 2 | 7.19 | **0.003** |
| Protein content | Temp | 1 | 0.32 | 0.578 | 1 | 11.6 | **0.002** |
| | Diet | 2 | 0.71 | 0.499 | 2 | 1.65 | 0.208 |
| | Temp*Diet | 2 | 2.51 | 0.098 | 2 | 1.41 | 0.259 |
| ETRmax | Temp | 1 | 28.16 | **<0.001** | 1 | 7.52 | 0.010 |
| | Diet | 2 | 63.68 | **<0.001** | 2 | 69.9 | **<0.001** |
| | Temp*Diet | 2 | 10.60 | **<0.001** | 2 | 12.8 | **<0.001** |

**Table S1**. Summary of two-way ANOVA for the physiological and biochemical parameters in *Pocillopora damicornis* and *Turbinaria reniformis* after thermal stress (Experiment 1 - Phase 2). Significant p values ($p \leq 0.05$) are in bold.

**Table S2.**

| Parameter | Effect | Phase 2 | | |
|---|---|---|---|---|
| | | *df* | *F* | *p* |
| Symbiont density | Temp | 1 | 5.24 | **0.031** |
| | Food | 2 | 1.36 | 0.274 |
| | Temp*Food | 2 | 11.9 | **<0.001** |
| Chlorophyll ($cm^2$) | Temp | 1 | 70.5 | **<0.001** |
| | Food | 2 | 2.55 | 0.077 |
| | Temp*Food | 2 | 5.23 | **0.0426** |
| Chlorophyll (cell) | Temp | 1 | 60.9 | **<0.001** |
| | Food | 2 | 4.23 | 0.064 |
| | Temp*Food | 2 | 1.31 | 0.102 |
| Protein content | Temp | 1 | 0.11 | 0.643 |
| | Food | 2 | 1.31 | 0.176 |
| | Temp*Food | 2 | 0.82 | 0.176 |
| ETRmax | Temp | 1 | 140.3 | **<0.001** |
| | Food | 2 | 34.3 | **<0.001** |
| | Temp*Food | 2 | 62.1 | **<0.001** |
| Catalase | Temp | 1 | 48.4 | **<0.001** |
| | Food | 2 | 12.7 | **<0.001** |
| | Temp*Food | 2 | 4.14 | **0.010** |
| GSH/GSSG | Temp | 1 | 10.6 | **0.001** |
| | Food | 2 | 4.22 | 0.140 |
| | Temp*Food | 2 | 4.05 | **0.015** |
| LPO (host) | Temp | 1 | 6.15 | **0.021** |
| | Food | 2 | 4.35 | **0.024** |
| | Temp*Food | 2 | 0.29 | 0.749 |
| LPO (symbiont) | Temp | 1 | 17.4 | **<0.001** |
| | Food | 2 | 1.48 | 0.246 |
| | Temp*Food | 2 | 12.8 | **<0.001** |
| TAC | Temp | 1 | 1.04 | **0.035** |
| | Food | 2 | 4.51 | **0.046** |
| | Temp*Food | 2 | 1.93 | 0.611 |

**Table S2**. Summary of two-factorial ANOVA for the physiological and biochemical parameters in *Stylophora pistillata* after thermal stress (Experiment 2 - Phase 2). Significant p values ($p \leq 0.05$) are in bold.

### Table S3.

| Parameter | Effect | Phase 3 | | |
|---|---|---|---|---|
| | | *df* | *F* | *p* |
| Symbiont density | Temp | 1 | 143.1 | **<0.001** |
| | Food | 2 | 11.3 | **<0.001** |
| | Temp*Food | 2 | 0.47 | 0.631 |
| Chlorophyll ($cm^2$) | Temp | 1 | 16.3 | **<0.001** |
| | Food | 2 | 4.95 | 0.066 |
| | Temp*Food | 2 | 1.43 | 0.355 |
| Chlorophyll (cell) | Temp | 1 | 231.7 | **<0.0001** |
| | Food | 2 | 4.40 | 0.177 |
| | Temp*Food | 2 | 4.76 | 0.198 |
| Protein content | Temp | 1 | 9.97 | **0.013** |
| | Food | 2 | 4.41 | **0.029** |
| | Temp*Food | 2 | 3.85 | **0.051** |
| ETRmax | Temp | 1 | 0.02 | **0.894** |
| | Food | 2 | 49.9 | **<0.001** |
| | Temp*Food | 2 | 64.4 | **<0.001** |
| ROS | Temp | 1 | 29.1 | **<0.001** |
| | Food | 2 | 7.03 | **0.006** |
| | Temp*Food | 2 | 0.96 | 0.588 |

**Table S3**. Summary of two-factorial ANOVA for the physiological and biochemical parameters in *Stylophora pistillata* after recovery (Experiment 2 - Phase 3). Significant p values ($p \leq 0.05$) are in bold.
